# Advanced methodology for bacterial colonization of 3D organotypic epidermal models: a gateway to long-term host-microbe interaction and intervention studies

**DOI:** 10.1101/2023.06.21.545853

**Authors:** Gijs Rikken, Luca D. Meesters, Patrick A.M. Jansen, Diana Rodijk-Olthuis, Ivonne M.J.J. van Vlijmen-Willems, Hanna Niehues, Peter Oláh, Bernhard Homey, Joost Schalkwijk, Patrick L.J.M. Zeeuwen, Ellen H. van den Bogaard

## Abstract

**Background:** Following descriptive studies on skin microbiota in health and disease, mechanistic studies on the interplay between skin and microbes are on the rise, for which experimental models are in great demand. Here, we present a novel methodology for microbial colonization of organotypic skin and analysis thereof.

**Results:** An inoculation device ensured a standardized application area on the *stratum corneum* and a homogenous distribution of bacteria, while preventing infection of the basolateral culture medium even during prolonged co-culture periods for up to two weeks at a specific culture temperature and humidity. Hereby, host-microbe interactions and antibiotic interventions could be studied, revealing diverse host responses to various skin-related bacteria and pathogens.

**Conclusions:** Our methodology is easily transferable to a wide variety of organotypic skin or mucosal models and different microbes at every cell culture facility at low costs. We envision that this study will kick-start skin microbiome studies using human organotypic skin cultures, providing a powerful alternative to experimental animal models in pre-clinical research.

## INTRODUCTION

The skin is a multi-faceted barrier organ that hosts a diversity of commensal microbial communities, composing the human skin microbiota. Over the past decade, we have witnessed a scientific breakthrough with respect to our knowledge and understanding of these microorganisms due to advances in sequencing technologies and the initiation of the human microbiome project [1]. Skin microbiome composition and diversity varies between body sites and individuals and is affected by environmental influences [2, 3]. The most abundant bacteria identified at the genus level are *Corynebacterium*, *Cutibacterium* and *Staphylococcus* [2, 4], along with the most common fungal commensal *Malassezia* [4–6]. These microbes play an important role in skin health by educating the immune system [7–9], preventing the colonization by pathogens [10, 11] and promoting skin barrier function [12, 13].

Alterations in skin microbiome composition, called dysbiosis, are nowadays associated with a plethora of skin conditions, such as atopic dermatitis (AD), psoriasis and acne [14–21]. Colonization and infection of the skin by *Staphylococcus aureus* (*S. aureus*) has been under investigation for decades [22, 23], but recent studies also suggest other *Staphylococcus* species like *S. epidermidis* [24] and *S. capitis* [25] to contribute to skin pathologies. The question remains whether dysbiosis is the cause or consequence of skin diseases and to what extent the microbiome can be leveraged as a therapeutic target [26–28]. Following initial descriptive studies on the skin microbiome [4, 29], investigative mechanistic studies using biologically relevant experimental models are of utmost importance to dissect the cause or contribution of microbial dysbiosis to health and disease [27, 30, 31].

Notwithstanding the importance and utility of widely used *in vivo*-animal models [32–34], the skin microbiome of rodents is significantly different from humans and the instability of the microbiome in laboratory animals is known to affect the experimental outcome [30]. Alternatively, human skin cell cultures (*e.g.,* keratinocyte monolayer cultures) allow investigations on the direct interaction between keratinocytes and microbes [35, 36]. Herein, co-cultures with live bacteria are restricted to be short-term as cell viability will be compromised upon the bacterial overgrowth within a few hours [37, 38]. Optionally, heat-killed bacteria, bacterial components or the bacterial culture supernatant can be used [39–41]. However, these do not mimic the actual colonization onto the protective *stratum corneum*, which acts as a physical barrier and filter for microbial metabolites [42]. Investigative studies on these metabolites and potential quorum sensing molecules [43, 44] that interact with bacterial or host cell receptors to activate signal transduction pathways [13, 45, 46], would benefit from models in which live bacteria are grown under biologically relevant culture conditions, such as a natural growth substrate (the *stratum corneum*) with a viable epidermis underneath.

Advanced organotypic skin models (either full-thickness skin or epidermal equivalents) have recently been used more often in host-microbe interaction studies. Next to bacterial infection models, microbial colonization is reported for a variety of skin-related bacteria and fungi. To summarize the current state-of-the-art, we provide a literature overview including experimental details and read-out parameters in Supplemental Table S1. These studies clearly indicate the utility of organotypic skin models for skin microbiome research, but also highlight a lack of standardization, relatively short co-culture periods of up to 24 hours, the high risk of basolateral culture infections and low assay throughput at high costs. Furthermore, the common use of standard cell culture conditions (37°C at a high relative humidity) in these microbial co-culture studies favors the growth of aerobic bacteria which will affect the bacterial diversity of *in vitro* cultured skin microbiome samples [47].

In an attempt to overcome these limitations, we here present a low cost and easy to use technical advance for microbial colonization of 3D human epidermal equivalents (HEEs). This may enable standardization of microbiome research using organotypic skin models and facilitate multi-parameter analytics from one single co-culture. Using this model system we provide proof-of-concept for differential host defense responses by skin commensals and pathogens, establish long-term co-culture periods up to two weeks and implement effective intervention studies by topical antibiotics.

## EXTENDED METHODS DESCRIPTION

## RESOURCES TABLE

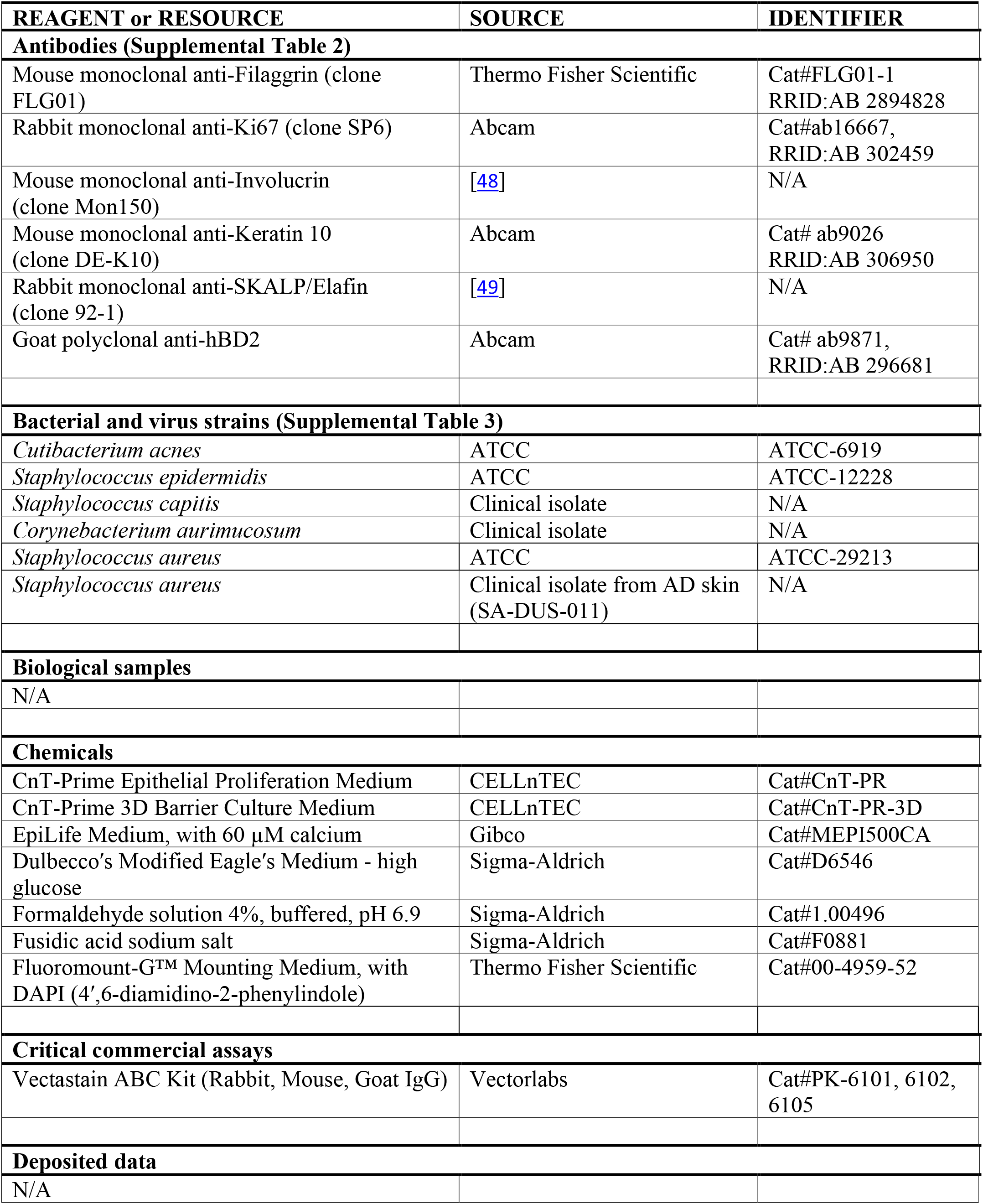
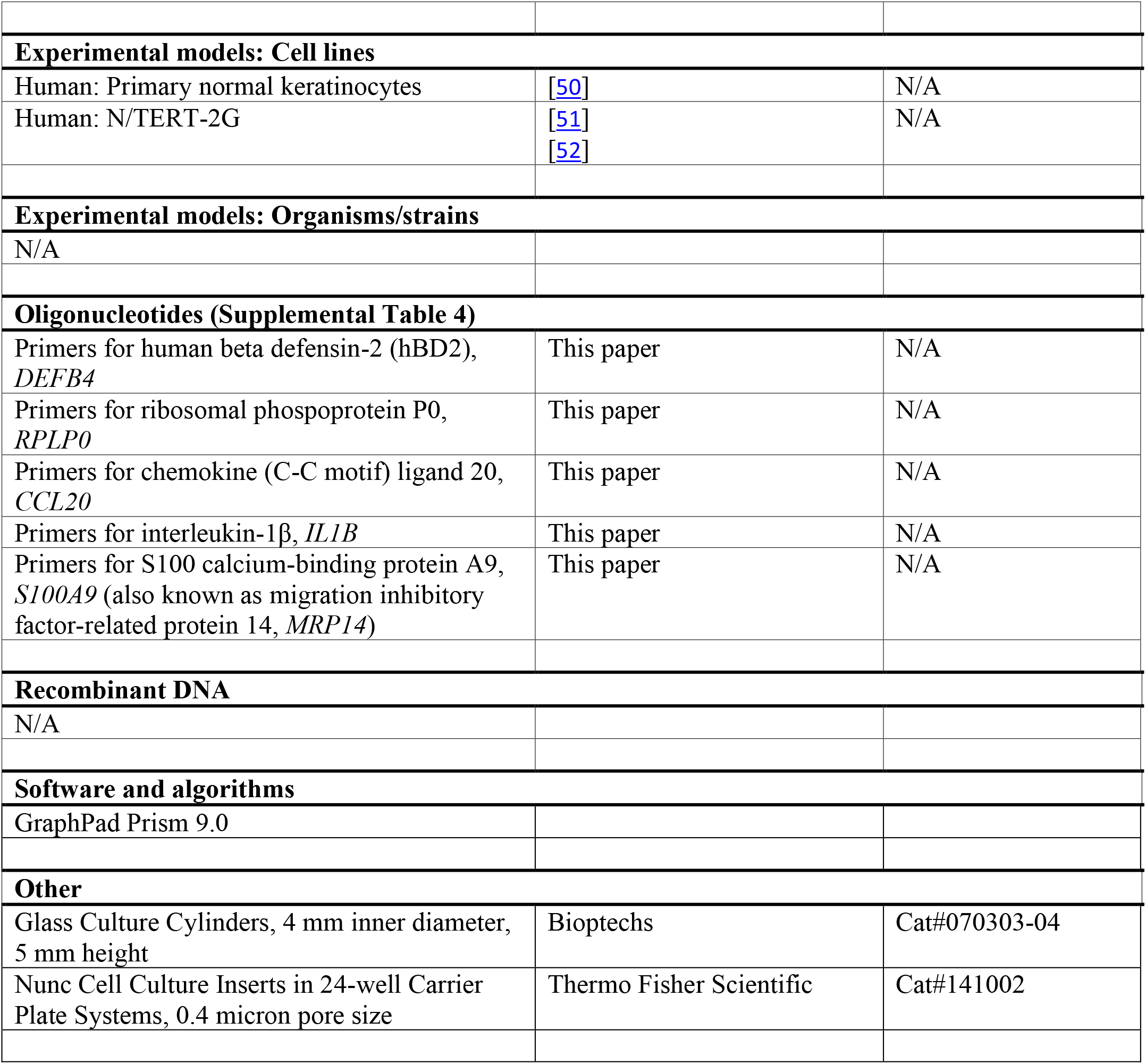

### CONTACT FOR REAGENT AND RESOURCE SHARING

Further information and requests for resources and reagents should be directed to and will be fulfilled by the lead contact, Ellen van den Bogaard.

## EXPERIMENTAL MODEL & METHOD DETAILS

### Primary keratinocyte isolation

Surplus human skin was obtained from plastic surgery (according to the principles of the Declaration of Helsinki). Human primary keratinocytes were isolated as previously described [50]. Briefly, 6 mm full-thickness biopsy punches of the freshly excised skin tissue were taken and placed into antibiotic/antimycotic medium for 4 hours at 4°C. Thereafter, 0.25% trypsin in phosphate buffered saline (PBS) was added and incubated overnight (o/n) at 4°C. Next, the enzymatic reaction was stopped by the addition of 10% (v/v) fetal bovine serum (GE Healthcare Life Sciences). A pair of tweezers was used to scrape the surface of the biopsy for harvesting of the keratinocytes. The keratinocytes were counted and seeded onto feeder cells at a density of 50.000 cells/cm^2^ in keratinocyte growth medium. The cells were harvested at 95% confluency with a final DMSO concentration of 10% and the cryovials were placed o/n into a freezing container at −70°C, after which the cells were stored in liquid nitrogen.

### 3D human epidermal equivalent (HEE) culture

HEEs were generated according to the protocols previously described (Rikken *et al*. 2020). Briefly, cell culture inserts (24-wells, 0.4 µm pore size filters; Thermo Fisher Scientific, Nunc) were coated with 150 µL of rat tail collagen in sterile cold PBS (100 µg/mL, BD Biosciences, Bedford, USA) at 4°C for 1 hour. Thereafter, excessive collagen solutions were carefully aspirated and the filters were washed with sterile cold PBS. Then, 150.000 primary human keratinocytes were seeded submerged in 150 µL CnT-prime medium (CELLnTEC, Bern, Switzerland). 900 µL of CnT-prime was added to the basolateral chamber, after which the cultures were incubated at 37°C and 5% CO_2_. After 48 hours, cultures were switched to 3D differentiation medium, which consists of 60% CnT-Prime 3D Barrier medium (CELLnTEC, Bern, Switzerland) and 40% High Glucose Dulbecco’s Modified Eagle’s Medium (DMEM, D6546, Sigma-Aldrich). 24 hours later, the HEEs were lifted to the air-liquid interface (ALI) using 1600 µL of 3D differentiation medium, which was refreshed every other day. The HEE culture schedule is depicted in Figure 1F (created with Adobe Illustrator, https://www.adobe.com/illustrator).

**Figure 1.**
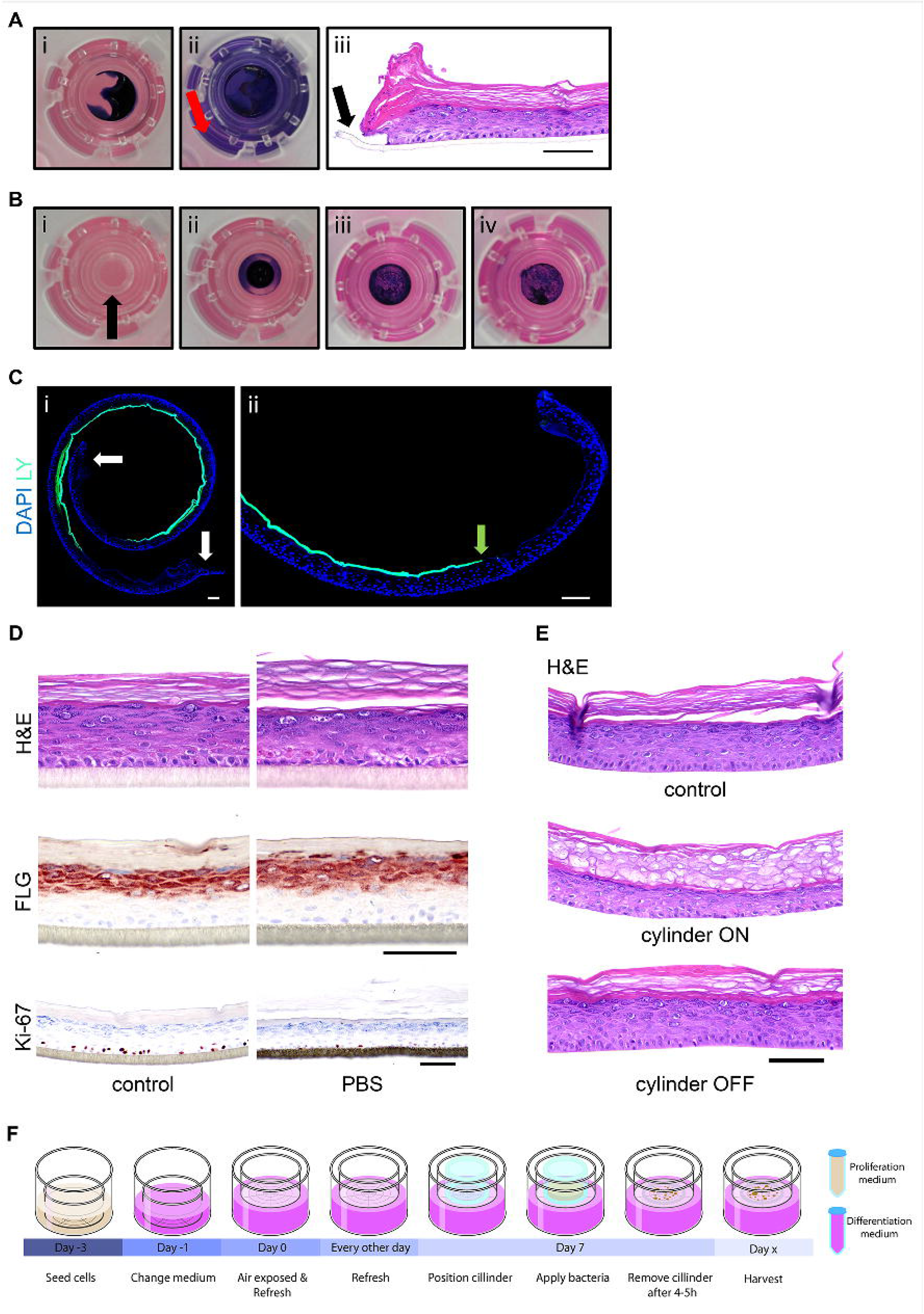
Validation of glass cylinder methodology. **(A)** i) 25 µL drop of trypan blue in PBS applied on top of the HEE, ii) the basolateral penetration of trypan blue after 4 hours of incubation (red arrow) and iii) H&E staining showing the open edges of the HEE (black arrow). **(B)** i) Glass cloning cylinder on top of the HEE indicated with the black arrow, ii) 25 µL of trypan blue in PBS was pipetted inside the cylinder, iii) the PBS was evaporated 4 hours later (in flow cabinet on heated plate at 37°C, without lid) and iv) the removal of the cylinder revealed a blue colonized circle without basolateral penetration. **(C)** Lucifer yellow (LY) added inside the glass cylinder and harvested after 2.5 hours of incubation. DAPI staining and fluorescent imaging (10x magnification) shows i) the distribution of LY onto the whole HEE and ii) clean edges. **(D)** H&E, Ki-67 and filaggrin (FLG) staining of HEE with a drop of PBS on top for 24 hours to analyze the morphological changes and protein expression patterns compared to the control. **(E)** Difference in morphology between the removal of the cylinder after PBS evaporation or leaving it on top of the HEE for 48 hours shown with an H&E staining. **(F)** Schematic overview of HEE culture and the topical application of bacteria using a glass cylinder. Scale bar = 100 µm.

For the N/TERT-2G cells, EpiLife medium (Gibco) or CnT-prime (CELLnTEC) was used (based on availability) for seeding the cells and during the first 48 hours of submerged culture. The N/TERT-2G HEEs were generated from N/TERT-2G keratinocytes at passage 3.

### Bacterial cultures

Bacterial strains (see Supplemental Table S2) were obtained from the Department of Medical Microbiology of the Radboud University Medical Center and the Department of Dermatology of the Heinrich-Heine-University in Düsseldorf (clinical isolate of AD skin, SA-DUS-011). *S. aureus*, *S. epidermidis*, *S. capitis* and *Corynebacterium aurimucosum* (*C. aurimucosum*) strains were grown o/n on Columbia agar with 5% sheep blood (Becton, Dickinson and Co.) under aerobic conditions at 37°C. Single colonies were used to inoculate cultures in 3 mL brain heart infusion (BHI) medium (Mediaproducts BV) in a 14 mL round bottom tube with snap cap (Cat#352057, Falcon, Corning) and incubated o/n at 37°C while shaking (225 rpm). Thereafter, bacterial cultures were diluted 100 times (30 µL in 3 mL BHI medium) and grown for another 2.5 hours in a shaking incubator to reach exponential growth. *Cutibacterium acnes* (*C. acnes*) was grown on Columbia agar with 5% sheep blood for 2 days at 37°C under anaerobic conditions (anaerobic jar system with an Oxoid Anaerogen 3.5L sachet (Cat#AN0035A, Thermo Fisher Scientific)), after which a single colony was picked and cultured o/n in 3 mL BHI medium at 37°C under anaerobic conditions. Thereafter, the bacteria were collected by centrifugation. The pellets containing the bacteria were washed twice in PBS and finally resuspended in PBS to reach the desired amount of colony forming units (CFU)/mL.

### Glass cylinder methodology for topical application of bacteria

After resuspension, the bacterial strains were topically applied on the *stratum corneum* of the organotypic cultures using a glass cloning cylinder (Cat#070303-04, Bioptechs, Pennsylvania, USA) with an outer diameter of 6 mm (inner diameter of 4 mm). Cylinders were first washed with soap followed by disinfection with 70% and 100% ethanol (air-dried in flow cabinet). The cylinder was placed on top of the HEE, with the raw surface facing downwards in the middle of the insert, using forceps, leaving approximately 1 mm space at the edge of the culture area. 25 µL of bacterial suspension (or PBS only) was carefully pipetted inside the cylinder. During 4-5 hours, the cultures were placed on a heated plate (37°C) in the flow cabinet (without the lid) to allow the surface to become dry again. Afterwards, the cylinder was carefully removed and additional supplementation of culture medium (approximately 100 µL) in the basolateral compartment was required before returning co-cultures to the incubator at 37°C and 5% CO_2_. A macroscopic view of the glass cylinder on top of the HEE is shown in Figure 1B, whereas a schematic overview of the HEE co-culture schedule is depicted in Figure 1F. During the co-culture experiments, samples of the culture medium were brought onto blood agar plates and incubated o/n at 37°C to check for sterility.

Depending on the experimental design, the bacteria were applied at different time points of the ALI (day 7, 8 and 11) and HEEs were harvested after 6 hours up to 13 days of co-culture. For the N/TERT-2G co-culture experiment, *S. aureus* ATCC 29213 was colonized at day 9 of the ALI.

To mimic the *in vivo* skin environment and to optimize co-culture conditions, HEEs inoculated with the SA-DUS-011 strain were also cultured at 32°C (at the start of colonization, up to 10 days) at low relative humidity. Of note, the culture medium in the basolateral chamber thereby evaporated faster requiring additional culture medium supplementation of 200 µL every day. Alternatively, the medium level could be increased with 500 µL to account for the evaporation and prevent the HEEs from running dry o/n.

The glass cylinder methodology was compared to a small droplet application (5 µL volume of bacterial suspension (SA-DUS-011 strain)) without the cylinder. The droplet was pipetted in the middle of the HEE (to minimize the risk of basolateral infections) and thereafter subjected to the same protocol as described above (37°C and 32°C).

### Topical application of antibiotics

Fusidic acid (FA, F0881, Sigma-Aldrich) was used as a narrow spectrum antibiotic known to combat *S. aureus* infections. Both *S. aureus* ATCC 29213 and the SA-DUS-011 strain were analyzed after the addition of FA in a concentration series. Immediately after the colonization of *S. aureus* (∼4 hours later, complete evaporation of PBS), 25 µL of FA (1% DMSO in water) was applied inside the same cylinder as used for the application of *S. aureus*. Again, the liquid was allowed to evaporate inside the flow cabinet (without lid on a heated plate, 37°C) and the cylinders were carefully removed afterwards. The HEEs with *S. aureus* ATCC 29213 were subjected to 1, 10 and 100 µg/mL FA, incubated at 37°C and 5% CO_2_ and harvested after 24 hours (technical triplicates).

For a prolonged HEE co-culture experiment with the SA-DUS-011 strain, FA (10 and 100 µg/mL) was applied every other day using the sterile glass cloning cylinder on top of the HEE. Co-cultures were incubated at 32°C (dry incubator) with 5% CO_2_ and harvested after 24 hours (technical triplicates) and 8 days (technical quadruplicates) of colonization.

## ANALYSIS METHOD DETAILS

### Multi-parameter end point analysis of organotypic co-cultures

The polycarbonate filter supporting the organotypic culture was gently pressed out of the transwell by placing it up-side-down and using a 8 mm biopsy punch (BP-80F, KAI Medical). A 6 mm biopsy punch was used to sample the area that had been covered by the glass cylinder. The bacterial colonization area was macroscopically visible to the naked eye, which allowed the precise excision using the biopsy punch. Of this 6 mm sample, a 3 mm biopsy was punched and fixed for 4 hours in 4% formalin for histological processing. The remainder of the sample was divided in two, with one part placed in 350 µL lysis buffer for total RNA isolation and the remainder in 250 µL PBS for CFU count, or in 500 µL PBS for microbial genomic DNA isolation for 16S rRNA gene sequencing. In summary, also depicted in the schematic image in Supplemental Figure S2B, samples were obtained for i) tissue morphology and/or protein expression ii) bacterial growth and iii) host gene expression from one single HEE to minimize batch effects and increase assay throughput.

### Immunohistochemistry

6 µm paraffin sections were stained with hematoxylin and eosin (Sigma-Aldrich) or mounted with DAPI (4′,6-diamidino-2-phenylindole) fluoromount-G (Thermo Fisher Scientific) after deparaffinization. For immunohistochemical analysis, sections were first blocked with 5% normal goat, rabbit or horse serum in PBS for 15 minutes and incubated with the primary antibody for 1 hour at room temperature or o/n at 4°C (Supplemental Table S3). Thereafter, the sections were washed in PBS and subsequently incubated with biotinylated secondary antibodies for 30 minutes. Next, sections were washed again in PBS and incubated with avidin-biotin complex (1:50 avidin, 1:50 biotin in 1% BSA/PBS (v/v)) (Vector laboratories) for 30 minutes. Protein expression was visualized by color change due to the peroxidase activity of 3-amino-9-ethylcarbazole (AEC). The tissue was counterstained with hematoxylin, after which the sections were mounted with glycerol gelatin (Sigma-Aldrich, Cat No. 1002946952).

### Keratinocyte RNA isolation and RT-qPCR analysis

RNA from the epidermal cells was isolated with the E.Z.N.A. Total RNA Kit I (OMEGA bio-tek) according to the manufacturer’s protocol. Isolated RNA was treated with DNaseI (Invitrogen) and used for cDNA synthesis using SuperScript IV VILO Master Mix (Invitrogen) and UltraScript 2.0 (PCR Biosystems) according to the manufacturer’s protocols. Subsequent real-time quantitative PCR (RT-qPCR) was performed using SYBR Green (Bio-Rad). qPCR primers were obtained from Biolegio (Nijmegen, The Netherlands) and depicted in Supplemental Table S4. Target gene expression levels were normalized using the house keeping gene human acidic ribosomal phosphoprotein P0 (*RPLP0*). The ΔΔCt method was used to calculate relative mRNA expression levels [53].

### Bacterial analysis

To isolate the bacteria from the organotypic co-cultures, the sample was homogenized/disintegrated in 250 µL PBS using a plastic micro pestle (Bel-Art, USA) in a 1.5 mL Eppendorf tube with convex bottom, by turning it around 10 times. Then, the suspension was completely homogenized using a needle (BD Microlance, 0.8 mm x 50 mm) and syringe (Henke-Ject, Tuberculin, 1 mL) by passing it 10 times. The homogenate was used to prepare a 10x dilution series and plated out on Columbia agar with 5% sheep blood. Plates were incubated at 37°C either o/n at aerobic conditions or for 2 days at anaerobic conditions. CFUs were counted and corrected for dilution and harvesting method, considering that only a part (3/8) of the co-culture was used for counting.

### Dye penetration assay

To determine the time point of *stratum corneum* formation allowing bacterial colonization, 25 µL of 1 mM lucifer yellow (LY, Sigma-Aldrich) was applied inside a glass cylinder on top of the HEEs at various time points of the ALI culture (day 5 till day 8) and incubated for 2.5 hours at 37°C. After routine formalin fixation and embedding in paraffin, 6 µm sections were counterstained and mounted using DAPI Fluoromount-G (Thermo Fisher Scientific). LY was visualized at excitation wavelength of 488 nm using the ZEISS Axiocam 305 mono and a 10x or 40x objective.

### Statistical analysis

Statistical analysis was performed using GraphPad Prism 9.0 (https://www.graphpad.com). Each HEE culture experiment includes technical replicates from a single keratinocyte donor, unless specified otherwise in the figure legend.

For the RT-qPCR gene expression analysis, the raw ΔCt values were used. An unpaired t-test was performed to determine statistical significance between two groups. Paired (biological replicates) and unpaired one-way analysis of variance (ANOVA) was used for comparison between multiple groups followed by Tukey’s multiple comparison post hoc test.

To determine statistical significance for the CFU count data, unpaired nonparametric one-sided Mann‒Whitney U test was used.

## DECLARATIONS

### Ethics approval and Consent to participate

The primary cells used in this study were obtained from surplus material from plastic surgeries according to the Declaration of Helsinki. Patients consent was documented in electronic patient records on the use of biological material that remained after treatment for scientific research (Code of Good Conduct). Bacterial isolates were obtained from non-invasive skin swabs patients after informed consent.

### Consent for publication

All authors have read and approved the manuscript prior to publication

## Availability of data and materials

Data or materials are available upon request by the senior author.

## Authors’ contributions

GR, PZ and EvdB designed the research. GR, LM and DR performed the cell and bacterial culture experiments and subsequent analysis, under supervision and expert advice from HN and PJ. IV assisted in the immunohistochemical analysis. GR and EvdB wrote the manuscript with feedback from PZ and JS. BH and PO provided the AD clinical isolate and advised on data interpretation. All authors read and approved the final manuscript.

## Acknowledgements

The authors thank undergraduate students Berber Maste, Priscilla Faas, Laura Edo Aceña, Blanca Gonzalez Melarde and Jaimy Klijnhout for technical assistance during their research internships which all contributed to the evolvement of the final model system. Danique van der Krieken assisted with the microbiological techniques, and Priscilla Faas created the schematic figures using Adobe Illustrator.

## Authors’ information (Optional)

Not applicable

## RESULTS

### The prerequisites for bacterial colonization of organotypic skin *in vitro*

For bacterial colonization of organotypic skin and the study of host-microbe interactions, prevention of cell culture infection is crucial. Like in native intact skin, the *stratum corneum* of organotypic skin models should form a barrier preventing bacteria from penetrating the epidermis. Therefore, the start of bacterial inoculation heavily depends on the correct *stratum corneum* formation of the organotypic HEEs to discriminate bacterial colonization from invasive infection. The first appearance of lipid-rich *stratum corneum* layers that marks the time point of inoculation can be easily visualized by a tracer molecule, lucifer yellow (LY). For all primary keratinocyte donors (N=8), LY was retained in the *stratum corneum* at day 7 of the air-liquid interface (ALI) culture, which was therefore considered as the starting point for bacterial colonization of HEEs in further experiments (Supplemental Figure S1A). To address the suitability of the HEE model for long-term bacterial co-culture studies, the lifespan of the HEEs was monitored. Expression patterns of the proliferation marker Ki-67, differentiation markers keratin 10 (K10) and filaggrin (FLG) and antimicrobial peptide (AMP) SKALP/elafin remained normal [54] for 25 days. The number of *stratum corneum* layers, however, increased due to lack of desquamation *in vitro* (Supplemental Figure S1B). After 30 days, a reduced number of epidermal layers and loss of the granular layer was seen (Supplemental Figure S1C). Therefore, the window of opportunity for studying host-microbe interactions or intervention strategies in the herein presented HEE model system was estimated being 18 days: from the start point of co-culture at day 7 of the ALI to maximally day 25.

### Glass cylinder methodology for standardized topical inoculation of HEEs

In our efforts to optimize the bacterial application method for inoculating HEEs (from small to larger bacterial suspension droplets or complete coverage of the HEE), we were challenged by the labor-intensiveness, lack in reproducibility of bacterial colonization, high inter-individual variation between researchers, detrimental effects on epidermal morphology and most importantly frequent immediate infections (<24 hours of co-culture) of the basolateral culture medium via the edges of the HEE. We therefore considered the utility of a glass cloning cylinder for topical application of the bacteria. The inert material minimally interacts with the bacteria or epidermis and allows easy sterilization. To quickly monitor the containment of liquid inside the cylinder at macroscopic level, we visualized the distribution of trypan blue on the HEE without and with the glass cylinder (Figure 1A-B, respectively). Microscopic analysis after LY application indicated an equal distribution over the *stratum corneum*, containment of liquid within the cylinder area and foremost clean edges of the HEE (Figure 1C).

Next, we investigated the effects of the glass cylinder and proposed vehicle (PBS) on the viability and structural integrity of the HEE. Prolonged immersion of organotypic epidermis is less desirable considering the detrimental effects on skin barrier formation and function [55, 56]. Indeed, covering HEEs with PBS for 24 hours changed the expression of markers for epidermal proliferation (Ki-67) and terminal differentiation (FLG) (Figure 1D). To reduce the time of liquid coverage of the *stratum corneum*, the cultures were placed on a laboratory hot plate (set at 37°C) without the lid of the transwell culture plate in the laminar flow hood to accelerate PBS evaporation. Thereby, the glass cylinder could be removed within 4-5 hours before returning the culture plates to the incubator. After careful morphological analysis (Figure 1E), this co-culture setup as depicted in Figure 1F was used as the basis for all further experiments.

### Inoculation of HEE with pathogens and skin commensals

For acquiring first proof-of-concept on our methodology, a bacterial suspension of the pathogen *S. aureus* (ATCC 29213, 10^4^ CFU in PBS) was added inside the glass cylinder, followed by a colonization period of 24 hours. Whole epidermal tissue analysis (8 mm biopsy punch) showed a homogenous distribution of the bacteria on the *stratum corneum* in the middle part, whilst keeping the edges of the HEEs free from bacteria (Supplemental Figure S2A). Next, we used one single HEE for multi-parameter readout analysis (Supplemental Figure S2B). After 24 hours of co-culture with two *S. aureus* strains (ATCC 29213 and a clinical isolate from an AD patient (SA-DUS-011)), CFU analysis indicated exponential bacterial growth reaching similar CFUs for both strains, with unaffected epidermal morphology (Figure 2A, Supplemental Figure S2C). Remarkably, marker gene expression analysis of AMPs (*DEFB4, S100A9* and *PI3*), revealed a strong induction after co-culture with SA-DUS-011 (Figure 2B). Also inflammatory mediators, here illustrated by *CCL20* and *IL1B*, were highly upregulated (Supplemental Figure S2D) in contrast to the laboratory ATCC strain.

**Figure 2.**
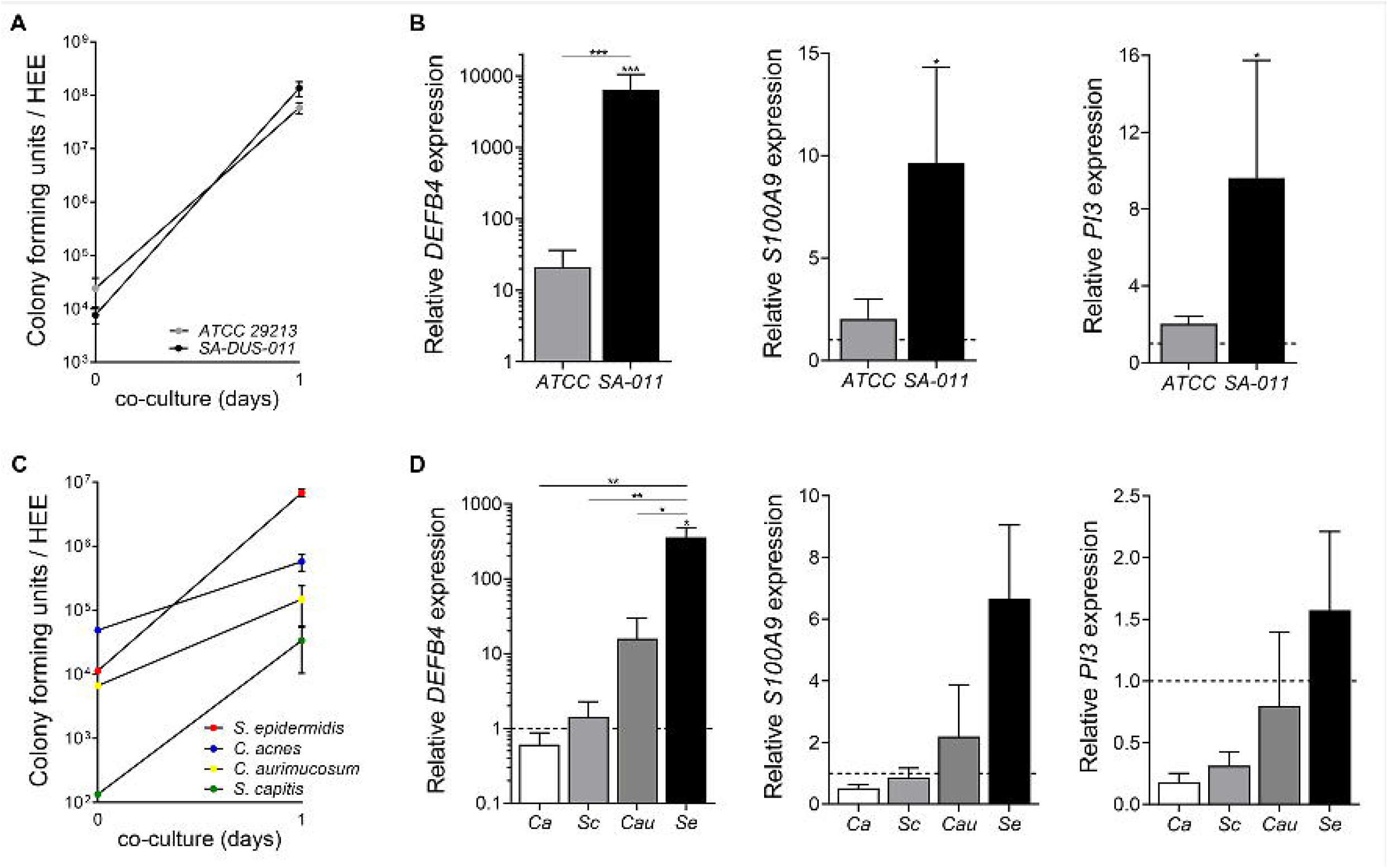
Colonization of HEEs with skin pathogens and commensals. **(A)** Growth and viability analysis by means of colony forming unit (CFU) count (input at 0 hours) (biological N*=*4) and **(B)** gene expression analysis of the antimicrobial peptides *DEFB4* (gene encoding hBD2), *S100A9* (MRP14) and *PI3* (SKALP/elafin) after 24 hours of co-culture with two *S. aureus* strains (ATCC 29213 and the clinical isolate SA-DUS-011) (biological N*=*4, all controls set at 1). **(C)** CFU count (input at 0 hours) (N*=*3) and **(D)** gene analysis of *DEFB4, S100A9* and *PI3* after 24 hours of co-culture with skin related bacteria (*S. epidermidis = Se, C. acnes = Ca, C. aurimucosum = Cau, S. capitis = Sc*) (N*=*3, control set at 1 (dashed line)). *p<0.05, **p<0.01. Mean ± SEM.

To study the capability of aerobic, aerotolerant or facultative anaerobic skin commensals to colonize HEEs, *S. epidermidis*, *S. capitis*, *C. aurimucosum* and *C. acnes* were co-cultured for 24 hours. CFU analysis indicated overall bacterial growth (Figure 2C), albeit at different growth rates between the tested strains (Supplemental Figure S2E). No differences were observed in the morphological appearance of the HEEs exposed to different bacterial strains (Supplemental Figure S2F), yet expression levels of host defense marker genes were significantly different, and mostly highly induced by *S. epidermidis* (Figure 2D, Supplemental Figure S2E and S2G). Importantly, no basolateral infections occurred during all HEE cultures as confirmed by plating culture medium onto blood agar plates.

### Prolonged co-culture of *S. aureus* ATCC 29213

Considering the favorable aerobic growth conditions for *Staphylococci* in HEE models and cell cultures in general, infections are expected upon long-term co-cultures if the glass cylinder does not effectively constrain the bacteria from leaking via the HEE edges, or when bacteria actively penetrate the *stratum corneum*. Being a commonly used human pathogenic strain, *S. aureus* ATCC 29213 was first selected for a prolonged two week co-culture period. *S. aureus* quickly reached a maximum of approximately 10^8^ CFU within 24 hours followed by a plateau phase during 13 days of co-culture (Figure 3A). The growth and survival of *S. aureus* on the HEE was irrespective of the start inoculum, reaching maximum levels between 10^7^ and 10^8^ CFU after 20 hours in all conditions (Supplemental Figure S3A). The epidermal morphology and protein marker expression for keratinocyte proliferation (Ki-67) and differentiation (FLG, involucrin (IVL)) of the HEEs co-cultured with *S. aureus* were comparable to control HEEs (Figure 3B, Supplemental Figure S3B). Induction of SKALP/elafin protein expression was observed after 24 hours of co-culture and remained stable over time (Figure 3C).

**Figure 3.**
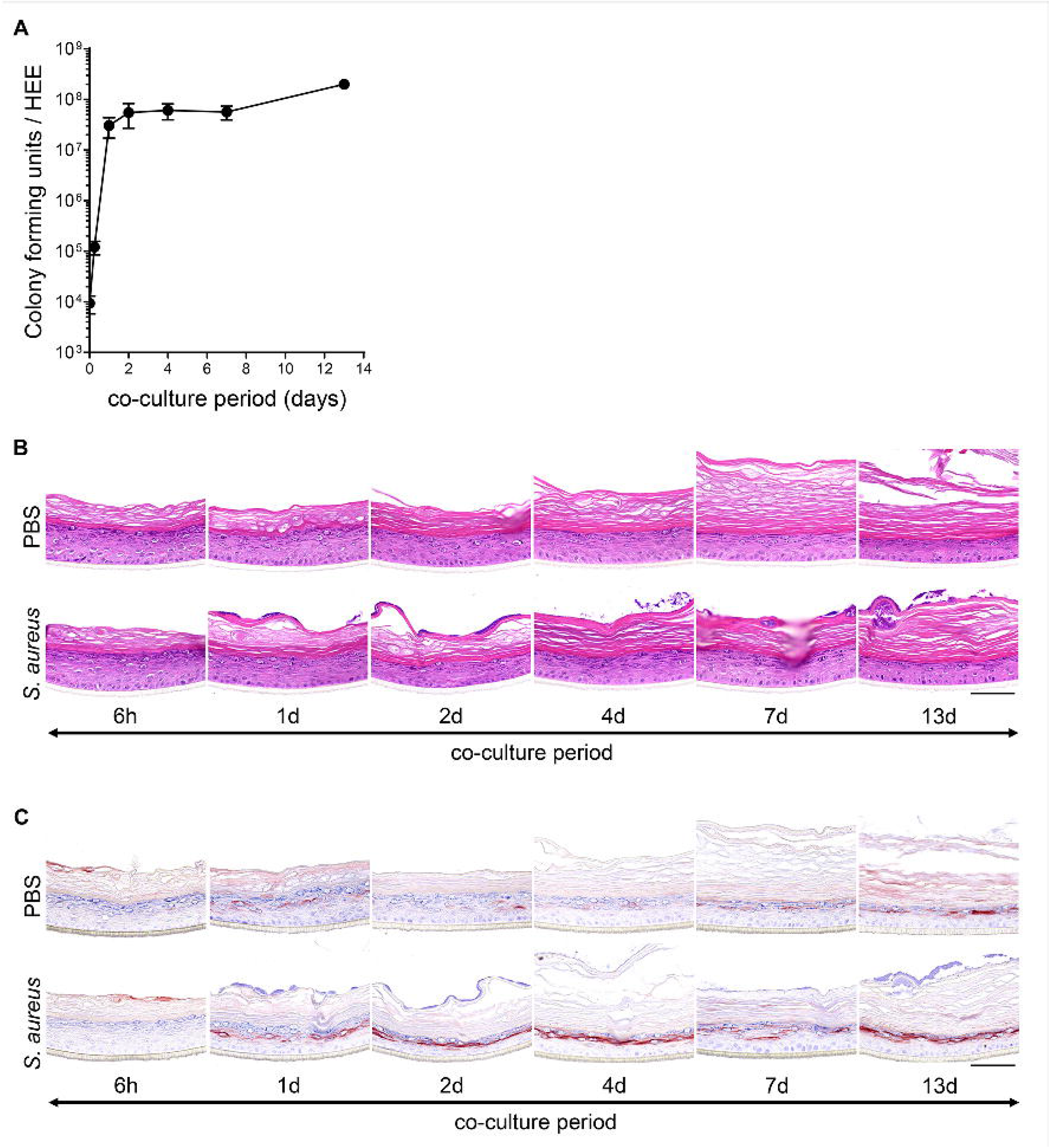
Prolonged co-culture analysis after *S. aureus* ATCC 29213 colonization. **(A)** Colony forming unit (CFU) analysis of HEEs inoculated with *S. aureus* ATCC 29213 and harvested at different time points of co-culture up to 13 days (input at day 0). All data points represent N*=*4 biological keratinocyte donor replicates, except for the 13 days co-culture (N*=*1). **(B)** H&E and **(C)** SKALP/elafin staining of the HEE donor co-cultured for 13 days with *S. aureus* and its vehicle (PBS). Scale bar = 100 µm.

Accumulating *stratum corneum* layers due to lack of desquamation *in vitro* (Figure 3B) could in principle hamper potential host-microbe interactions at later stages of the co-culture period. However, *stratum corneum* thickness did not influence bacterial growth and viability (Supplemental Figure S3C), nor did it hamper the induction of SKALP/elafin (Supplemental Figure S3D) when applying *S. aureus* at later stages of the ALI (day 11). Considering the popularity of the immortalized N/TERT keratinocytes in skin science as an alternative cell source for primary keratinocytes, we generated HEEs from the N/TERT-2G cell line which resulted in similar colonization rates as observed for primary keratinocytes (Supplemental Figure S4A-B). Again, in all experiments, no infections occurred during the short-term co-culture period.

### Epidermal infections after prolonged colonization by *S. epidermidis* and *S. aureus*

Commensal bacteria like *S. epidermidis* can become opportunistic pathogens causing skin infections [57] and may induce AD-like disease at high abundances [58]. Considering the strong host defense response we observed already after 24 hours of HEE colonization (Figure 2D, Supplemental Figure S2G), we evaluated the effects of a more prolonged co-culture with *S. epidermidis.* Epidermal infections occurred within 96 hours, even at a minimal input inoculum of 10^2^ CFU. Structural damage of the epidermis, including loss of the granular layer, parakeratosis and reduced epidermal layers was observed (Figure 4A). Strong induction of hBD2 and SKALP/elafin protein expression after 96 hours (Figure 4B) was subsequently accompanied by the confirmed presence of bacteria in the culture medium. Of note, no basolateral infections were observed after 24 hours, rejecting the hypothesis of initial infection via leaky HEE edges.

**Figure 4.**
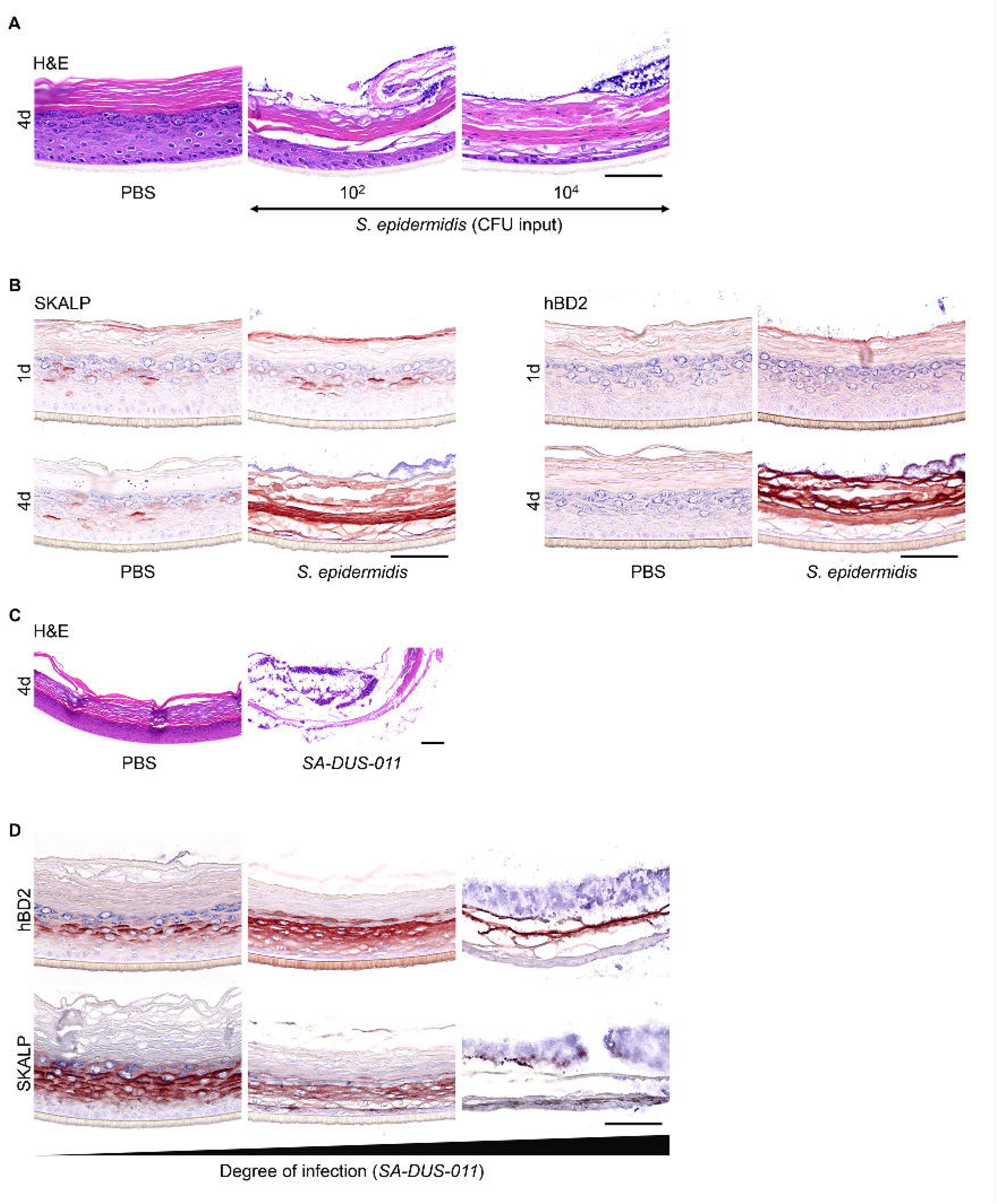
Epidermal infections caused by *S. epidermidis* and *S. aureus*. **(A)** *S. epidermidis* (10^2^ and 10^4^ CFU input) caused epidermal infections within 96 hours of co-culture, visualized with H&E staining that revealed the structural damage and loss of granular layer compared to the control HEE (PBS). **(B)** Immunostainings of the AMPs SKALP/elafin and hBD2 showed induction of protein expression in case of an epidermal infection. **(C)** H&E staining of HEE colonized with the *S. aureus* clinical isolate SA-DUS-011 (10^4^ CFU input) for 96 hours compared to the control HEE (PBS). **(D)** HEEs inoculated with SA-DUS-011, harvested at different time points of infection and stained for the AMPs SKALP/elafin and hBD2. All HEEs had multiple visible large yellow colonies on top of the *stratum corneum*. Only the culture medium of the first HEE was not infected yet, analyzed with a blood agar plate and o/n incubation at 37°C. Scale bar = 100 µm.

Since the *S. aureus* clinical isolate (SA-DUS-011) also showed strong induction of host defense gene expression at 24 hours, we also prolonged this co-culture, resulting in basolateral cell culture infections within 96 hours (Figure 4C). Prior to bacterial growth in the basolateral compartment, yellow colonies typical for *S. aureus* were macroscopically visible on the HEE surface after 48 hours. Harvesting the SA-DUS-011 HEEs at different time points indicated various degrees of infection by upregulated AMP expression (hDB2 and SKALP) at the start of infection followed by structural damage to the epidermis (Figure 4D). Similar results were obtained using N/TERT HEEs. Herein, epidermal infections were seen in 5/6 replicates after 72 hours with concomitant upregulation of *DEFB4* (Supplemental Figure S4C). The induction of antimicrobial proteins upon microbial co-culture may thus be considered as an indicator of an epidermal infection *in vitro*, even when the epidermal morphology is still unaffected and basolateral culture medium shows no signs of infection.

### Bacterial infections related to culture conditions

To address the influence of potential experimental artefacts (*e.g.*, *stratum corneum* defects) from the cylinder application, the glass cylinder methodology was head-to-head compared with the application of a small volume of SA-DUS-011 in the middle of the HEEs [59–61]. In addition, to better mimic the natural growth conditions of bacteria on skin, physiologically relevant co-culture conditions (32°C, dry atmosphere; cold and dry) were compared to the traditional cell culture conditions (37°C, high humidity; warm and humid).

The large bacterial surface area in the cylinder in warm and humid conditions conferred significantly higher CFU count and relative growth than the droplet area and reached similar CFU counts as in previous experiments (10^7^-10^8^ CFU) (Figure 5A-B). At cold and dry conditions, a maximum CFU of 10^6^ per HEE was reached at both the droplet and cylinder application method, albeit the number of HEEs that became infected significantly differed between both application methods (Figure 5C). Briefly, the smaller droplet area delayed infection onset in a warm and humid environment by at least 4 days, but could not prevent all HEEs becoming infected within 7 days of co-culture. Dry and cold conditions delayed infection onset using the cylinder and even prevented infections in 80% of HEEs with a small (droplet) application area.

**Figure 5.**
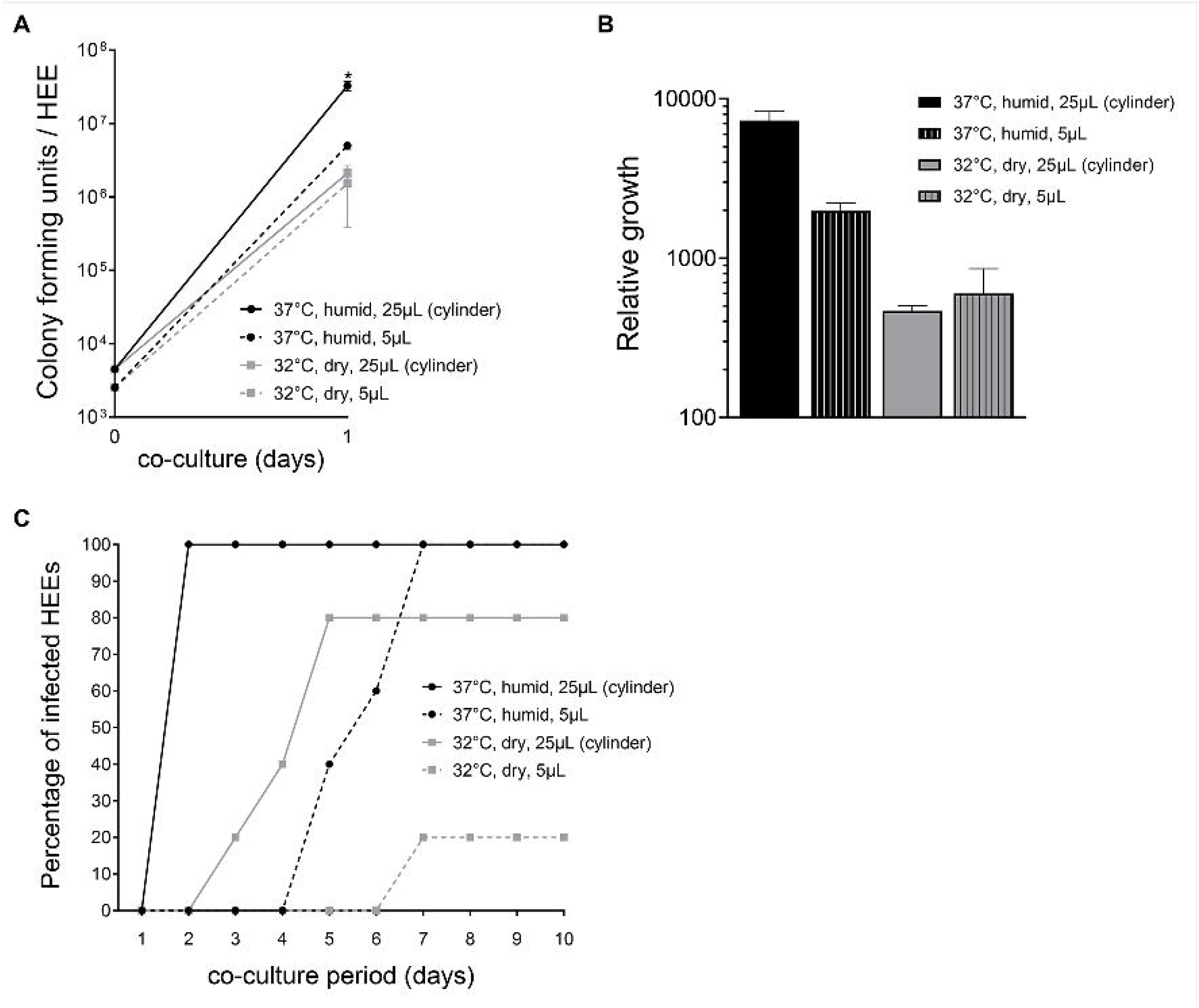
Bacterial infections using different culture conditions. **(A)** Colony forming unit (CFU) analysis and **(B)** relative growth of the *S. aureus* clinical isolate SA-DUS-011 after 24 hours of colonization applying four different methods (glass cylinder methodology (25 µL) *versus* small droplet (5 µL) and 37°C (humid) *versus* 32°C (dry)) (N*=*3 per method) (input at 0 hours), *p<0.05 (Mann-Whitney U test, CFU outcome of 37°C glass cylinder method compared to the other methods). **(C)** Percentage of infected HEEs (N*=*5 per method), co-cultured for up to 10 days with SA-DUS-011 applied using the four different methods.

To further dissect the influence of temperature versus humidity on bacterial growth and infection rate, HEEs were also cultured at 32°C in a humid environment. After 48 hours, SA-DUS-011 caused epidermal infections in all HEEs that were incubated in humid conditions (of note: the infections started earlier at 37°C compared to 32°C). In dry and cold conditions, only 3 out of 8 HEEs became infected.

### Topical antibiotic inhibits the growth of *S. aureus*

Next to more fundamental studies on skin host-microbe interactions, organotypic 3D skin microbiome models could be of importance for research and development of pre-, pro-and antibiotics to modulate the skin microbiome for therapeutic purposes. We implemented the cylinder methodology for the topical application of antibiotics using readout parameters for both host and microbe. Fusidic acid (FA) is used in clinical practice for the treatment of *Staphylococci* skin infections and herein chosen as a proof-of-principle intervention.

Inhibition of *S. aureus* ATCC 29213 growth was observed in a dose dependent manner after a single dose of FA was added inside the cylinder directly after the initiation of *S. aureus* colonization, indicating the bacteriostatic effect of FA (Figure 6A). In the morphological analysis, the lower amount of *S. aureus* colonies on top of the *stratum corneum* relate to the effective FA treatment. At the effective FA concentrations of 10 and 100 µg/mL, no morphological changes of the HEE were observed (Figure 6B). Based on the aforementioned optimal co-culture conditions, FA efficacy was tested (10 and 100 µg/mL) on the *S. aureus* clinical isolate SA-DUS-011 using the glass cylinder and culturing in a cold and dry environment up to 8 days. At day 1, CFU analysis showed a strong reduction of *S. aureus* (Figure 6C) indicative of the effective bacteriostatic effects of FA (bacteria were not completely killed, resulting in 10^5^ CFU on day 8 upon FA treatment every other day). During the following 7 days, 50% of the untreated *S. aureus*-colonized HEEs became infected after 4 days. The remainder of the untreated *S. aureus*-colonized HEEs that were harvested at day 8 showed severe epidermal damage (Figure 6D) with high CFU counts (Figure 6C) indicative of epidermal infections. FA treatment not only limited the bacterial growth, but also completely prevented infections and epidermal damage caused by *S. aureus* in HEEs.

**Figure 6.**
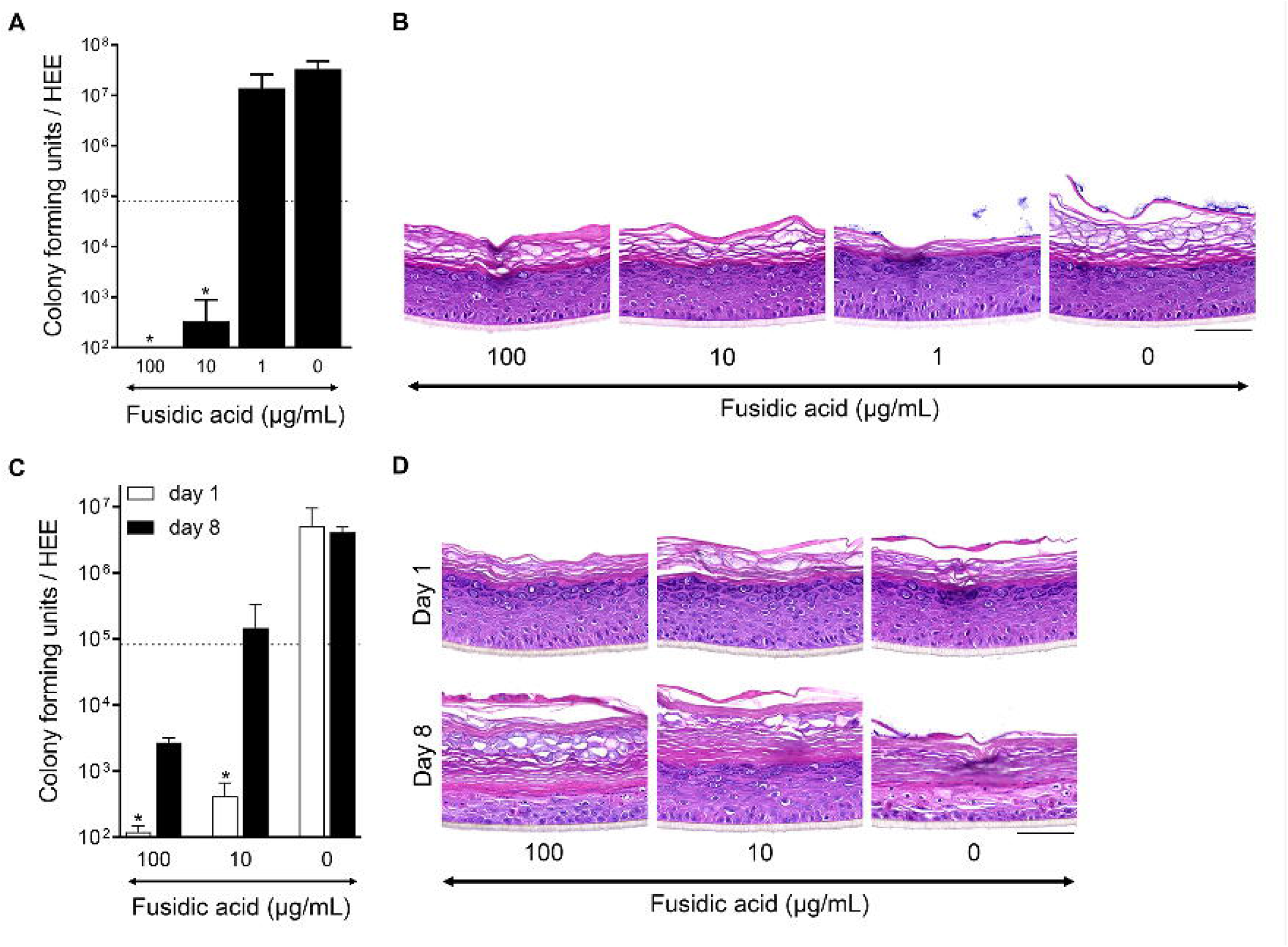
Fusidic acid inhibits the growth of *S. aureus* on HEEs. **(A)** Dose inhibiting effect via colony forming unit (CFU analysis) of HEEs topically applied with fusidic acid (FA, 1-10-100 µg/mL, 1% DMSO in water (0 µg/mL, vehicle)) 4 hours after *S. aureus* ATCC 29213 colonization (dotted line: amount of CFUs at start of treatment) and harvested 24 hours later (N*=*3), and **(B)** H&E staining thereof. **(C)** CFU analysis on day 1 (N*=*3 per treatment) and day 8 (0 µg/mL (N*=*2) and 10-100 µg/mL (N=4)), and **(D)** H&E staining of HEEs colonized with the *S. aureus* clinical isolate SA-DUS-011 subjected to the FA treatment protocol (10 and 100 µg/mL). HEEs were analyzed with a prolonged co-culture up to 8 days to study epidermal infections; 50% (2 out of 4) *S. aureus* HEEs infected after 96 hours (FA applied at day 0, 2, 4 and 6) (of note, co-cultured at 32°C (dry)). *p<0.05 (Mann-Whitney U test, CFU outcome of fusidic acid dosages compared to the vehicle (0 µg/mL)). Mean ± SEM. Scale bar = 100 µm.

## DISCUSSION

We here present a technical advance for the topical bacterial inoculation of a 3D human epidermal equivalent (HEE) with a minimal risk of basolateral infections, whilst allowing *in vitro* studies on infectious virulent strains. This methodology using glass cylinders will be easily transferable to a wide variety of advanced organotypic skin [62], [63] or mucosal models [64], would be amenable for the application of diverse microbiota (bacteria [65, 66], viruses [67–69] and fungi [70, 71]) and can be used in every cell culture facility considering the various sizes and commercial availability. We were able to increase the assay throughput by the large bacterial exposure area and thus obtaining multiple samples for various endpoint analysis from one single HEE.

We generated both a colonization and infection model based on the single strain exposure of a fully developed epidermal model. While other bacterial co-culture models to date induce an infection by making a wound [61, 63, 72, 73], we here showed that the *S. aureus* clinical isolate (SA-DUS-011) caused epidermal infections after colonizing an intact skin. Albeit similar growth rates and a high CFU output (10^7^-10^8^), the *S. aureus* strain ATCC 29213 did not infect the HEE within two weeks of co-culture nor did it induce the expression levels of any of the host defense markers, indicating a strain specific effect. Therefore, screening of various skin related bacterial species and using more than one strain per bacterium, ideally isolated from individual patients or volunteers, followed by whole genome sequencing [47], could relate virulence factors to the clinical features of the patient and host-microbe responses *in vitro*.

While here we present the model characteristics using single bacterial strains, the ultimate goal would be the application of whole skin microbiome samples or pre-designed microbial communities, as used in experimental animal models [13]. Yet, *in vitro* cell culture conditions have been shown to affect the stability of the commensal communities, skewing towards a dominance of aerobic bacteria after the co-culture period [47] and 16S or shotgun sequencing only includes information on relative abundancies whilst lacking information on bacterial viability. Methods to exclude bacterial DNA from dead cells, like propidium monoazide (PMA) [74], may provide a solution but require a labor-intense multi-step protocol and will be difficult to validate for the correct dosing of complex bacterial mixtures to avoid killing of microbes due to treatment.

The major advantage of a glass cylinder is the large colonization surface, allowing the collection of multiple samples, that we called “*multiple parameter endpoint analysis*”. A small droplet, as commonly used, prevents infection of the basolateral chamber, but will require multiple transwell inserts, large experimental setups or cell culture formats (6-12well), [59, 62, 65, 66, 75–77]. Others completely cover the cell culture surface with bacterial suspension, but this requires immediate analysis or removal of non-adherent bacteria [71, 78–81]. Furthermore, when the set-up of experiments require multiple treatment steps of the equivalents, the cylinder provides a defined area wherein treatments can be applied after each other by equally distributed evaporation of the solutions, as we here showed for fusidic acid. This antibiotic prevented infections and maintained the epidermal morphology for at least 8 days of treatment, which is a novel finding compared to other antibiotic organotypic models [59, 78, 80, 82]. Although we found that the glass cylinder does accelerate the start of epidermal infections, a small droplet application also resulted in infections. Therefore, we value the utility of the glass cylinder and changed the culture environmental conditions (32°C in a dry atmosphere) to delay the onset of infections and maximize the co-culture period and window of opportunity for interventions. By changing the cell culture environmental conditions and varying the application area of bacteria we leverage the opportunity to either study skin infection or colonization. Interestingly, we observed that under dry culture conditions, co-cultures located in the middle of the culture plate infected earlier than those in the outer rows, presumably due to higher humidity in the middle of the culture plate. Hence, only controlling the humidity in the cell culture incubator is not sufficient to fully standardize environmental conditions within the culture plate.

Modulation of microbiome composition and its effects might also be accomplished by changing host factors. We here showed that the use of the N/TERT-2G immortalized keratinocyte cell line is a suitable alternative for microbial colonization of HEEs since the epidermal structure is similar to that of primary keratinocytes [51]. In addition, it is the preferred cell type for genome editing and the use of a cell line instead of primary cells will reduce the biological variation. For example, knockdown of the differentiation protein filaggrin (*FLG*) showed increased colonization of *S. aureus* on top of the organotypic N/TERT model [79]. This correlation between FLG and microbial colonization is also observed *in vivo* for *S. aureus* [83, 84]. In addition, specific commensal species are underrepresented on *FLG*-deficient skin showing a reduction of gram positive anaerobic cocci [37], that appear to harbor important AMP-inducing capabilities [41]. Furthermore, continued efforts in the optimization of culture conditions and protocols to better mimic the *in vitro* skin barrier to that of native skin [85, 86] will also affect the interaction between microbes and epidermal keratinocytes in organotypic model systems and as such, it will remain a challenge to compare results obtained between various models. Detailed information on the model characteristics (morphology, skin barrier function, cell sources, culture medium, microbial strain selection) are pre-requisites for studies that aim to investigate cell-host-microbe interactions in organotypic skin models.

In conclusion, our developed model system allows for easy utilization of organotypic human epidermal models for investigative skin microbiome research. This opens avenues into the application of more complex microbial cultures, the evaluation of specific pathogens in genotype-defined organotypic human skin models, and the screening of pre-, pro-or antibiotic treatments therein.

---

**Supplemental Figure S1.**
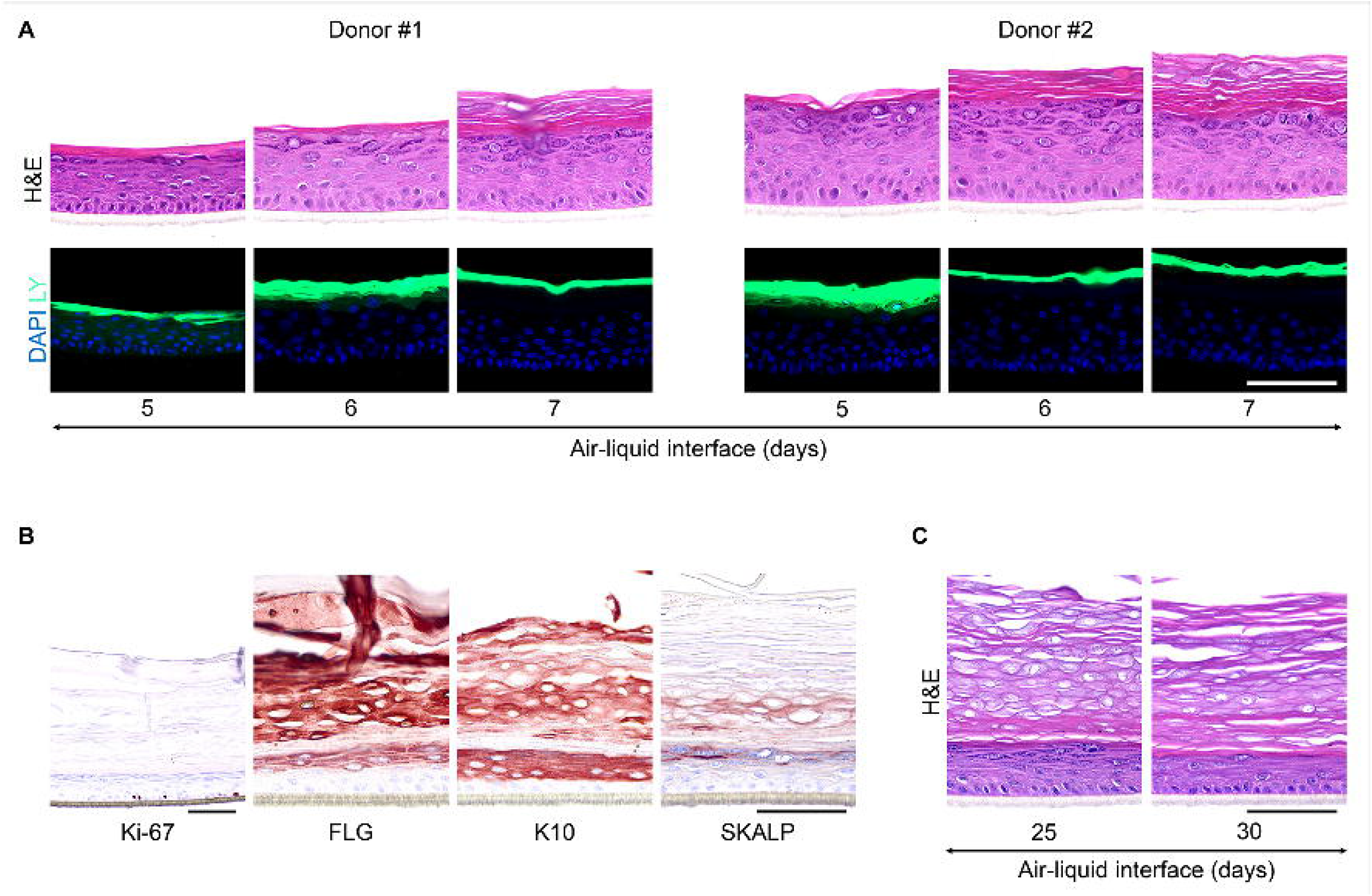
*Stratum corneum* formation and lifespan of HEEs. **(A)** H&E and DAPI staining of two HEE donors that were topically applied with LY for 2.5 hours on different days of the air-liquid interface (ALI) to evaluate *stratum corneum* penetration (images represent eight biological keratinocyte donors). **(B)** Protein expression of the proliferation marker Ki-67, differentiation markers filaggrin (FLG) and keratin 10 (K10) and the AMP SKALP/elafin of a HEE at day 25 of the ALI. **(C)** H&E staining of HEEs harvested at day 25 and 30 of the ALI to investigate the lifespan of the culture. Scale bar = 100 µm.

**Supplemental Figure S2.**
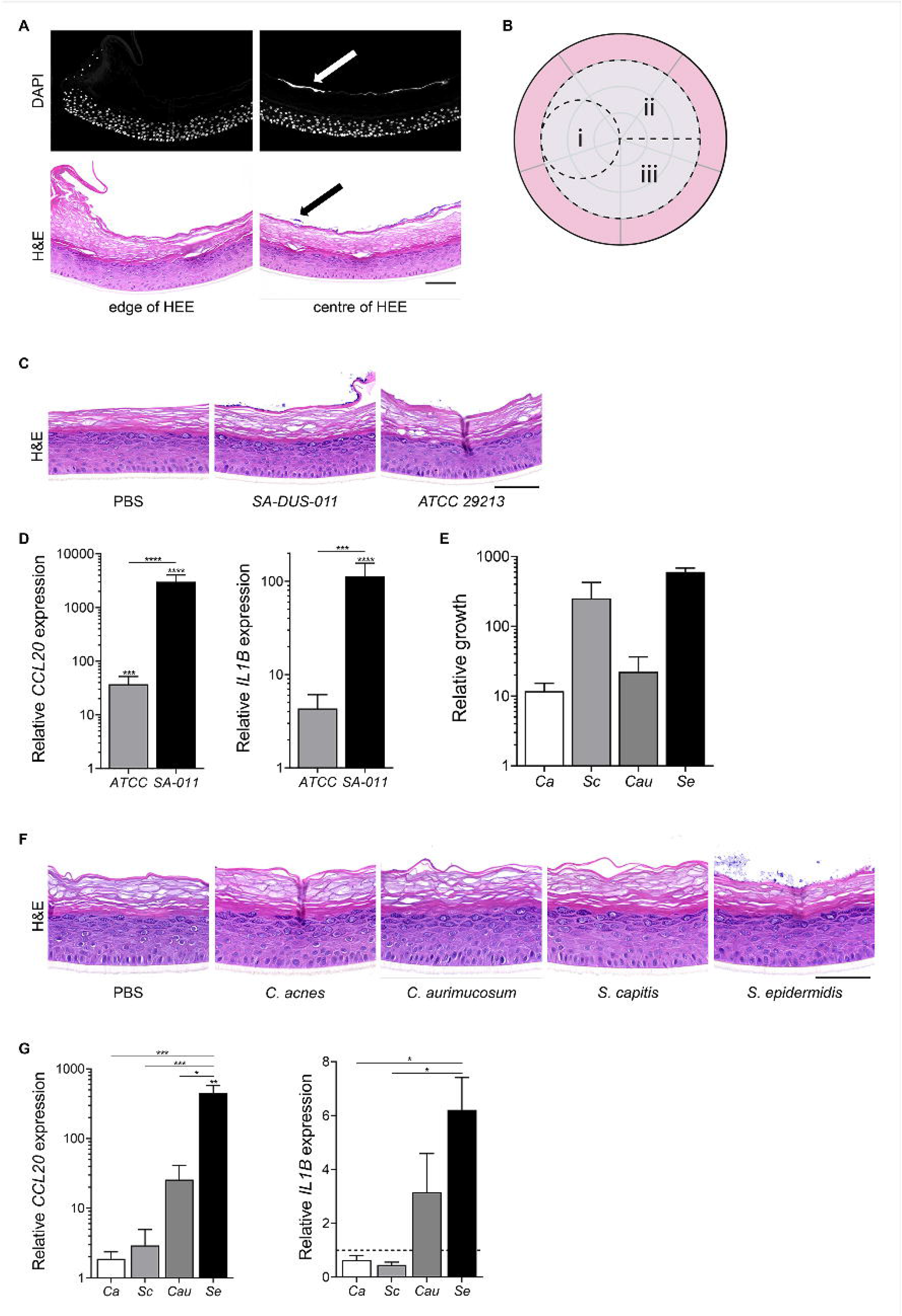
Multi-parameter endpoint analysis, bacterial colonization, growth and host defense response. **(A)** DAPI (white nuclei and colonies (white arrow)) and H&E (colonies indicated with black arrow) staining of HEE co-cultured for 24 hours with 10^4^ colony forming units (CFU) of *S. aureus* ATCC 29213 to visualize bacterial colonization and clean edges of the HEE. **(B)** Multi-parameter analysis for i) morphology and/or protein expression, ii) host gene expression and iii) bacterial growth. **(C)** H&E staining and **(D)** inflammatory gene expression (*CCL20* and *IL1B*) of HEEs colonized with *S. aureus* ATCC 29213 and the *S. aureus* clinical isolate SA-DUS-011 for 24 hours to analyze epidermal morphology (biological N*=*4, controls set at 1). **(E)** Logarithmic growth, **(F)** H&E staining and **(G)** inflammatory gene expression (*CCL20* and *IL1B*) after 24 hours of co-culture with skin related bacteria (*S. epidermidis = Se, C. acnes = Ca, C. aurimucosum = Cau, S. capitis = Sc)* (N*=*3, control set at 1). *p<0.05, ***p<0.001. Mean ± SEM. Scale bar = 100 µm.

**Supplemental Figure S3.**
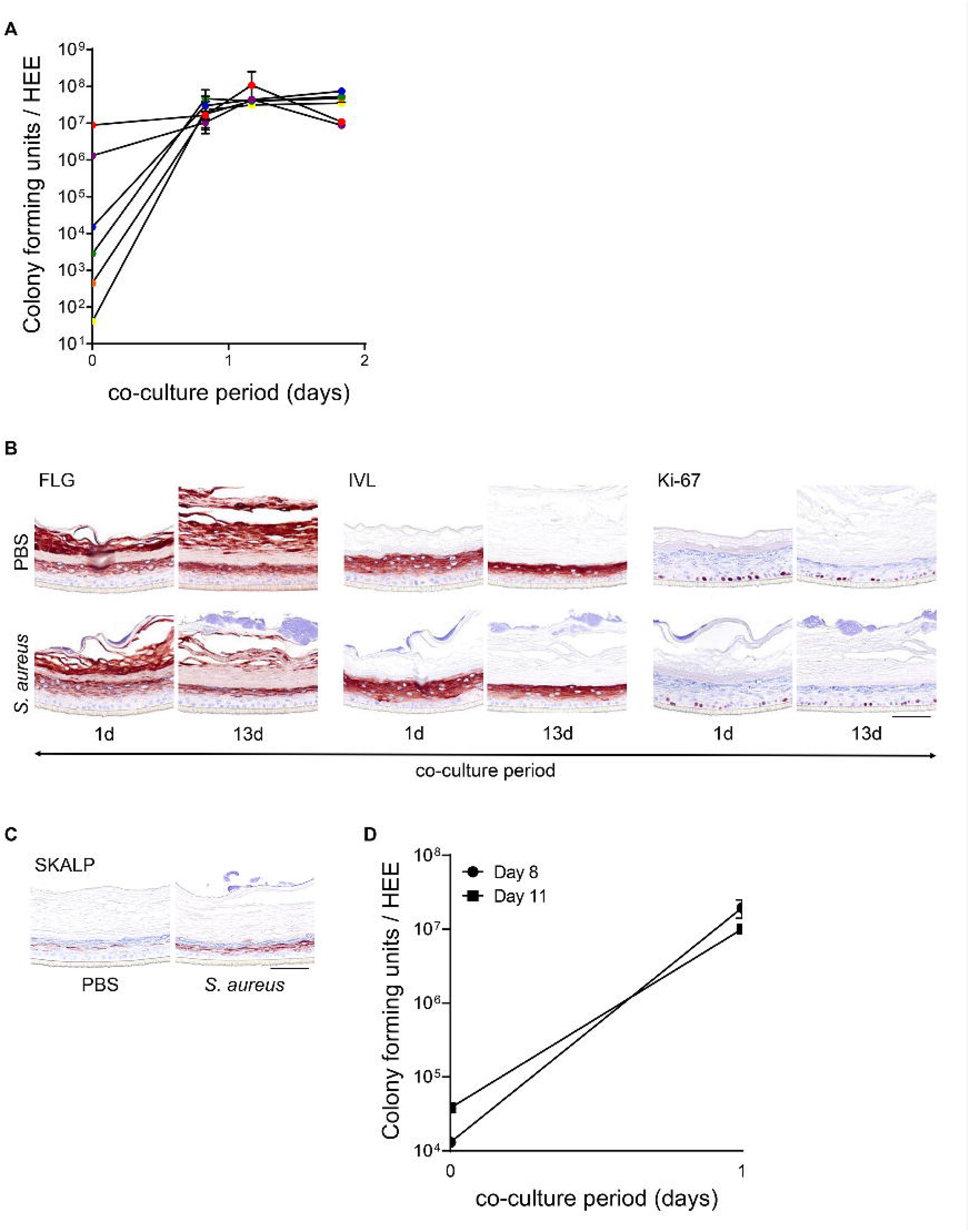
Inoculum and *stratum corneum* thickness do not influence growth of *S. aureus* ATCC 29213. **(A)** Colony forming unit (CFU) count of HEEs inoculated with a concentration series (10^1^, 10^2^, 10^3^, 10^4^, 10^6^ and 10^7^ CFU) of *S. aureus* and harvested after 20, 28 and 44 hours of co-culture (N*=*2). **(B)** Normal epidermal protein expression after *S. aureus* colonization up to 13 days compared to the control HEE (PBS) shown with the proliferation marker Ki-67 and the differentiation markers filaggrin (FLG) and involucrin (IVL). **(C)** CFU analysis of *S. aureus* colonized at day 8 and day 11 (thick layer of *stratum corneum*) of the air-liquid interface (ALI) for 24 hours (biological N*=*5, input at day 0). **(D)** SKALP/elafin protein expression of HEE inoculated with *S. aureus* at day 11 of the ALI (thick layer of *stratum corneum*) in comparison with the control HEE (PBS) and co-cultured for 24 hours. Images represent N*=*5 biological keratinocyte donors. Scale bar = 100 µm.

**Supplemental Figure S4.**
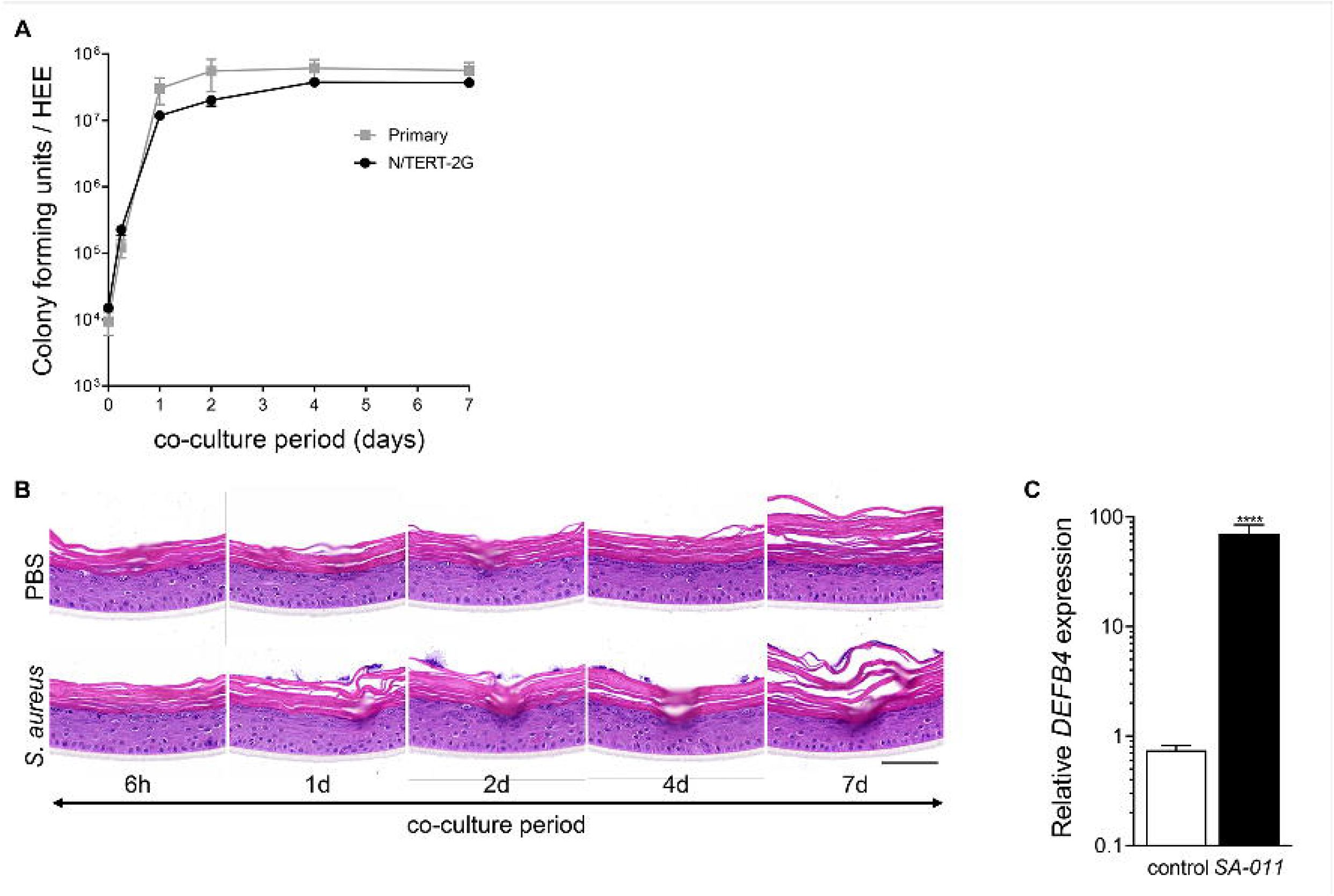
HEEs generated with immortalized N/TERT cells and colonized with *S. aureus* strains. **(A)** Colony forming unit (CFU) analysis of N/TERT HEEs colonized with *S. aureus* ATCC 29213 and harvested after different time points of co-culture up to 7 days (each data point N*=*3), in comparison with primary human keratinocytes (grey line, biological N*=*4) and **(B)** H&E staining thereof. **(C)** Gene expression analysis of the antimicrobial peptide *DEFB4* after 72 hours of co-culture with the *S. aureus* clinical isolate (SA-DUS-011) (N=6). ****p<0.0001. Mean ± SEM. Scale bar = 100 µm.

**Supplemental Table S1:**
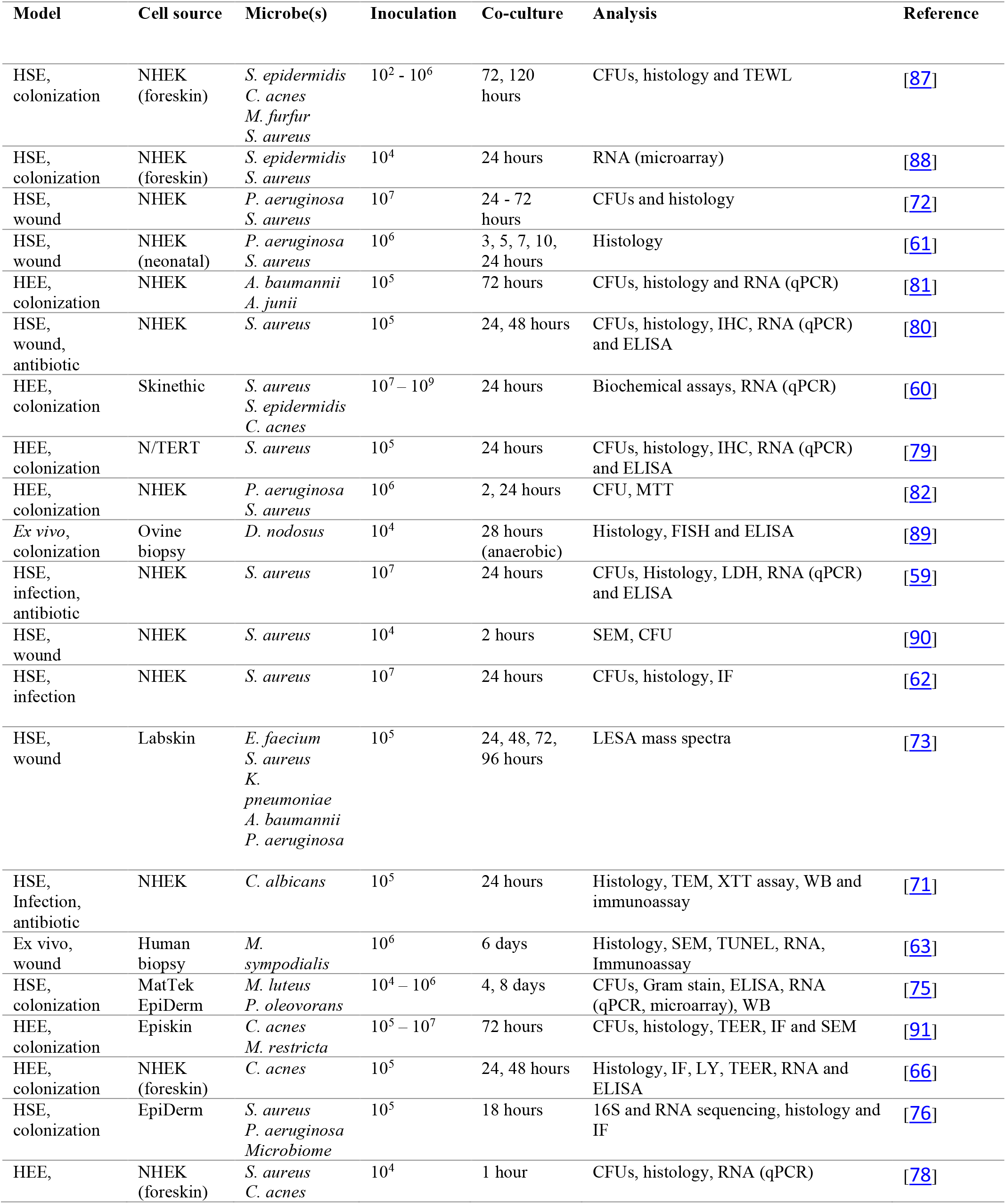
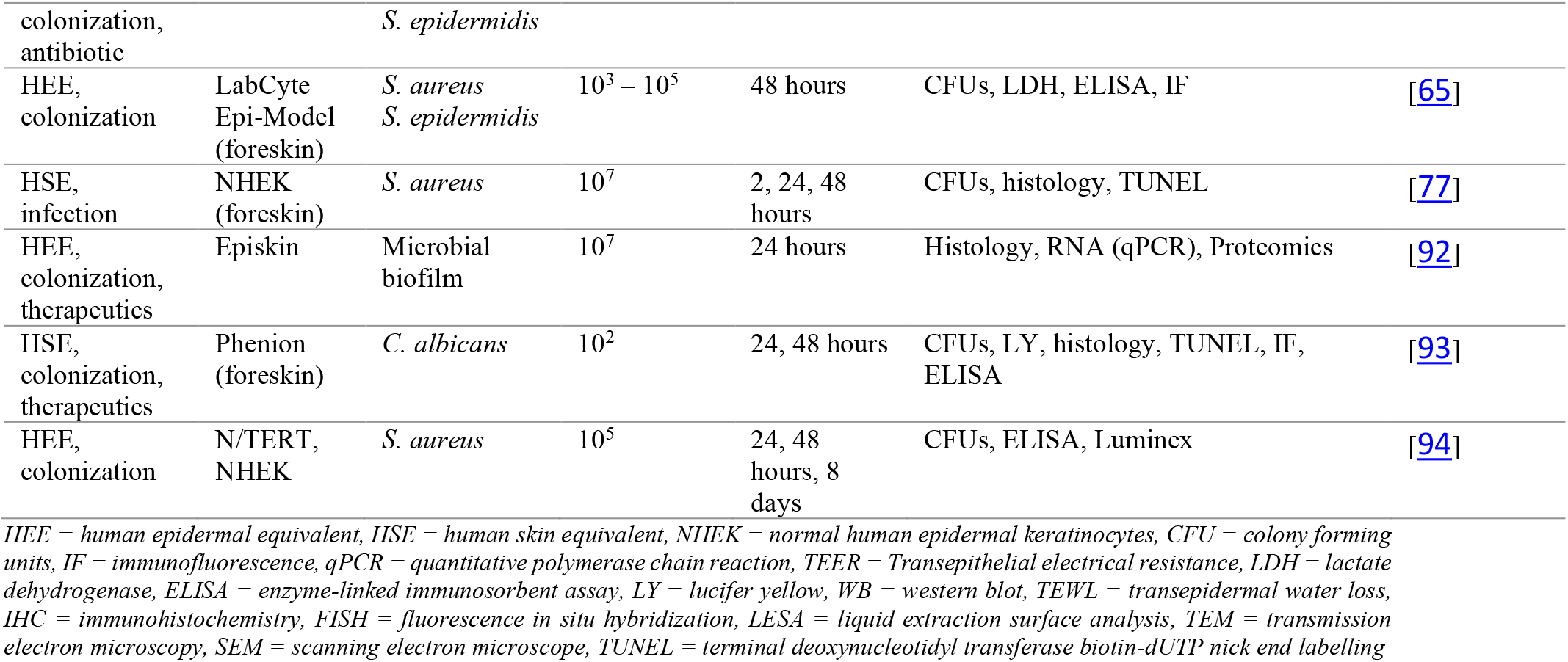
Studies that used 3D organotypic skin models to investigate bacterial colonization, infection and host-microbe interactions.

**Supplemental Table S2.**
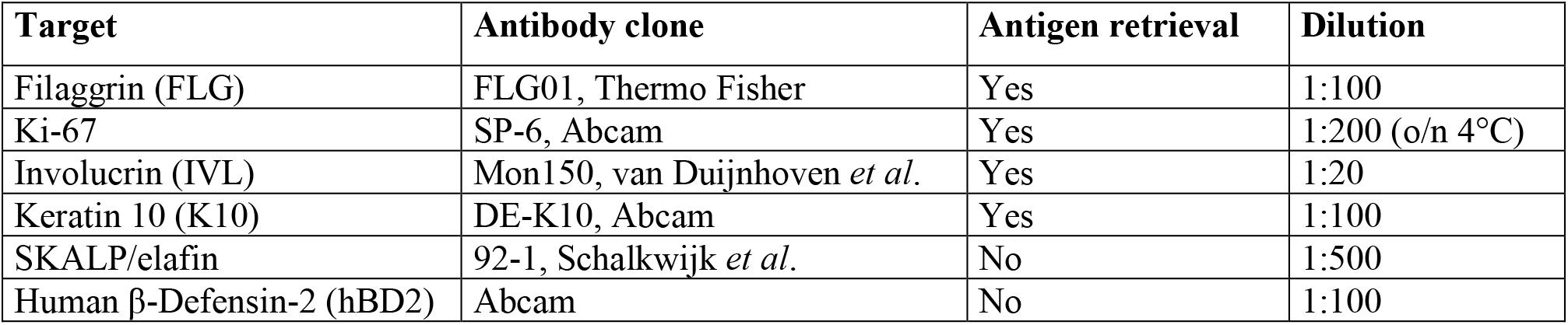
Antibodies used for immunohistochemistry

**Supplemental Table S3.** Bacterial strains

| Strain | Identification number | Gram | Growth conditions |
| --- | --- | --- | --- |
| *Cutibacterium acnes* | ATCC-6919 | positive | anaerobe |
| *Staphylococcus epidermidis* | ATCC-12228 | positive | aerobe |
| *Staphylococcus capitis* | Clinical isolate | positive | aerobe |
| *Corynebacterium aurimucosum* | Clinical isolate | positive | aerobe |
| *Staphylococcus aureus* | ATCC-29213 | positive | aerobe |
| *Staphylococcus aureus* | Clinical isolate | positive | aerobe |

**Supplemental Table S4.**
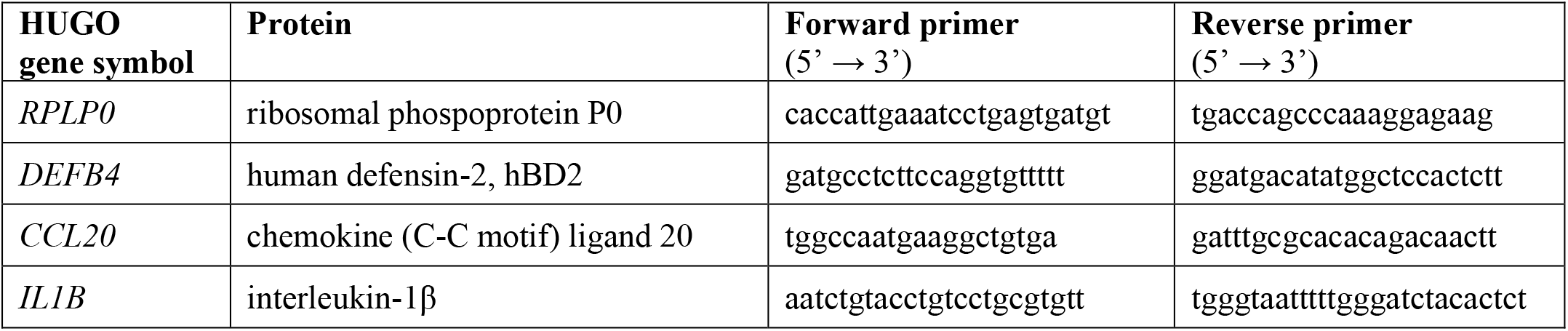
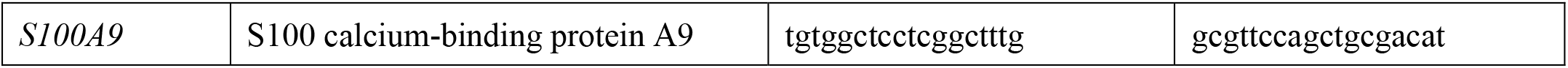
Primers for qPCR

